# Robust Mouse Tracking in Complex Environments using Neural Networks

**DOI:** 10.1101/336685

**Authors:** Brian Q. Geuther, Sean P. Deats, Kai J. Fox, Steve A. Murray, Robert E. Braun, Jacqueline K. White, Elissa J. Chesler, Cathleen M. Lutz, Vivek Kumar

## Abstract

The ability to track animals accurately is critical for behavioral experiments. For video-based assays, this is often accomplished by manipulating environmental conditions to increase contrast between the animal and the background, in order to achieve proper foreground/background detection (segmentation). However, as behavioral paradigms become more sophisticated with ethologically relevant environments, the approach of modifying environmental conditions offers diminishing returns, particularly for scalable experiments. Currently, there is a need for methods to monitor behaviors over long periods of time, under dynamic environmental conditions, and in animals that are genetically and behaviorally heterogeneous. To address this need, we developed a state-of-the-art neural network-based tracker for mice, using modern machine vision techniques. We test three different neural network architectures to determine their performance on genetically diverse mice under varying environmental conditions. We find that an encoder-decoder segmentation neural network achieves high accuracy and speed with minimal training data. Furthermore, we provide a labeling interface, labeled training data, tuned hyperparameters, and a pre-trained network for the mouse behavior and neuroscience communities. This general-purpose neural network tracker can be easily extended to other experimental paradigms and even to other animals, through transfer learning, thus providing a robust, generalizable solution for biobehavioral research.

## Main

Behavior is primarily an output of the nervous system in response to internal or external stimuli. It is hierarchical, dynamic, and high dimensional, and is generally simplified for analysis^1,2^. For instance, the rich locomotor movement performed by a mouse that is captured in video is routinely abstracted to either a simple point, a center of mass, or an ellipse for analysis. In order to do this well with current methods, the experimental environment is simplified to obtain optimal contrast between the mouse and background for proper segmentation. Segmentation, a form of background subtraction, classifies pixels belonging to mice from background in video and enables these high level abstractions to be mathematically calculated. During mouse experimental assays, the arena background color is often changed depending on the animal’s coat color, potentially altering the behavior itself^3-5^. Making such changes comes at a cost, as current video tracking technologies cannot be applied in complex and dynamic environments or with genetically heterogeneous animals without a high level of user involvement, making both long term experiements and large experiments unfeasible. As neuroscience and behavior moves into an era of big behavioral data^2^ and computational ethology^6^, current tracking methods are inadequate and improved methods are necessary that enable tracking animals in semi-natural and dynamic environments over long periods of time. To address this shortfall, we developed a robust scalable method of mouse tracking in an open field using modern convolutional neural network architecture. Our trained neural network is capable of tracking all commonly used strains of mice—including mice with different coat colors, body shapes, and behaviors—under multiple experimental conditions without any user-involved adjustment of tracking parameters. Thus we present a scalable and robust solution that allows tracking in diverse experimental conditions.

We first used existing tracking methods to track 59 different mouse strains in multiple environments, and found them inadequate for our large-scale strain survey experiment (1,845 videos, 1,691 hours). Specifically, we tracked all the videos in this experiment using Ctrax^7^, a modern open-source tracking software package that uses background subtraction and blob detection heuristics, and LimeLight (Actimetrics, Wilmette, IL), a commercially available tracking software package that uses a proprietary tracking algorithm. Ctrax abstracts a mouse on a per frame basis to five metrics: major and minor axis, x and y location of center of the mouse, and the direction of the animal^7^. It utilizes the MOG2 background subtraction model, whereby the software estimates both the mean and variation of the background of the video for use in background subtraction. Ctrax uses the shape of the predicted foreground to fit ellipses. LimeLight uses a single key-frame background model for segmentation and detection. Once a mouse is detected using LimeLight, this software package abstracts the mouse to a center of mass using a proprietary algorithm.

Our strain survey experiment includes videos of mice with different genetic backgrounds causing expression of different coat colors including black, agouti, albino, grey, brown, nude, and piebald (Fig. 1a, columns 1, 2, 3 and 4). We tracked all animals in the same open field apparatus, which had a white background; this yielded good results for darker mice (black and agouti mice), but poor results for lighter-colored (albino and grey mice) or piebald mice (Fig. 1a, columns 1, 2, 3 and 4, Supplementary Video 1). Examples of ideal and actual tracking frames are shown for the various coat colors (Fig. 1a, row 3 and 4 respectively).

**Figure 1:**
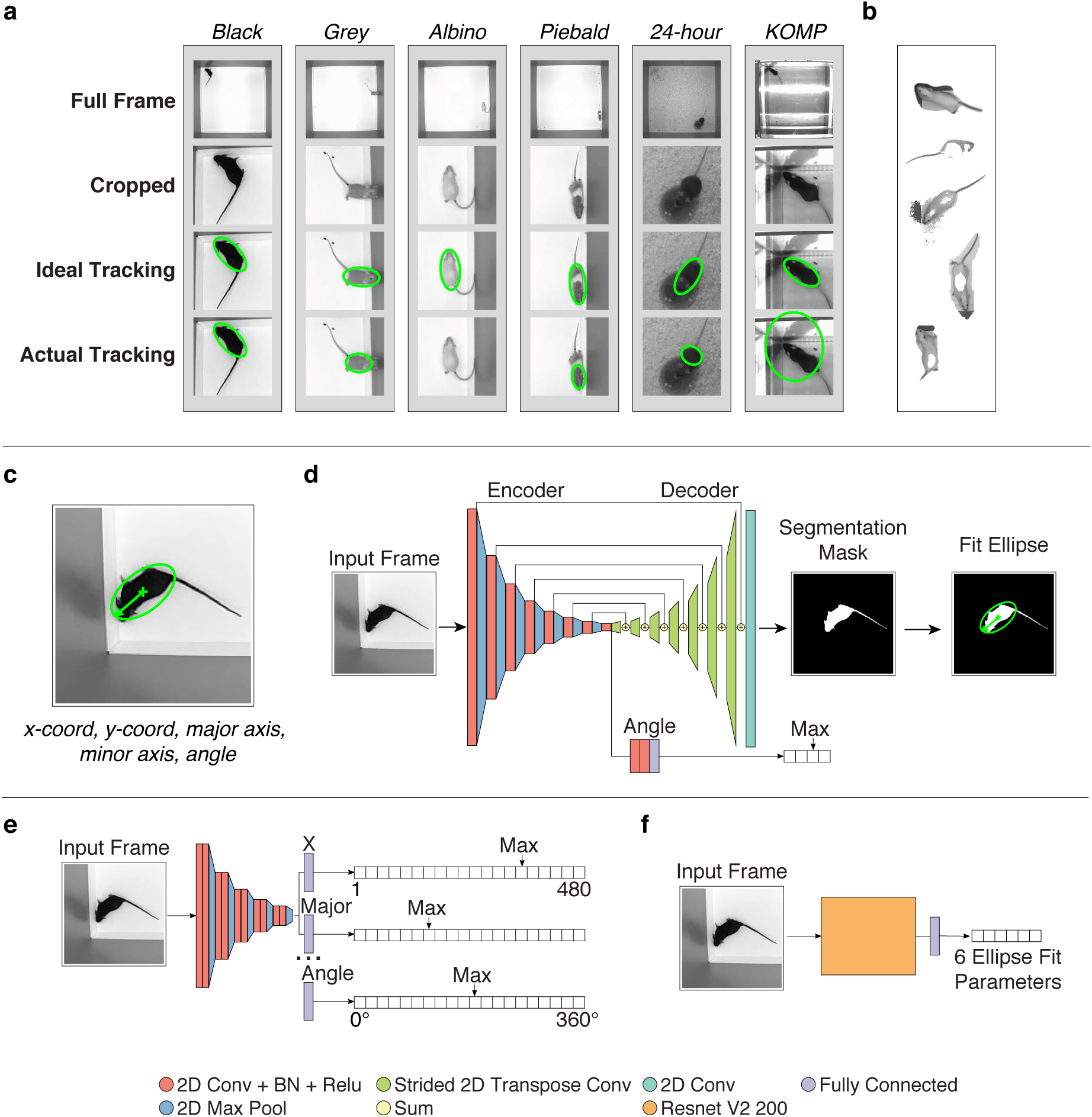
(**a**) A representation of the environments analyzed by existing approaches. A black mouse in a white open field achieves high foreground-background contrast, and therefore actual tracking closely matches the ideal (column 1). Grey mice are visually similar to the grey-colored arena walls and therefore often have their noses, which are grey, removed while rearing on walls (column 2). Albino mice are visually very similar to the white arena floor and are frequently not found during tracking (column 3). Piebald mice are broken in half by the tracking software due to their patterned coat color (column 4). Placing a food cup, that is visually similar to the mouse, into the arena causes tracking issues when the mouse climbs on top of the food cup (column 5). Arenas with reflective surfaces also produce errors with tracking algorithms (column 6). (**b**) We identified the cause of bad tracking as poor segmentation. However, testing a variety of difficult frames with multiple background subtraction algorithms from the background subtraction library, we did not resolve this segmentation issue. From top to bottom the background subtraction algorithms shown are: SuBSENSE, Adaptive Median, Adaptive Background Learning, MultiCue BGS, and LOBSTER. (**c**) Our objective tracking takes the form of an ellipse description of a mouse. For clarity, we show a cropped frame as input into the networks, whereas the actual input is an unmarked full frame. (**d**) The structure of the segmentation network architecture functions similarly to classical tracking approaches in which the network predicts the segmentation mask for the mouse and then fits an ellipse to the predicted mask. (**e**) The structure of the binned classification network architecture predicts a probability distribution of the value for each ellipse-fit parameter, represented by the table where a max value is selected. Only three parameters of the six ellipse-fit parameters are visually shown (X = center x-location, Major = major axis length, Angle = direction of the mouse’s nose). (**f**) The structure of the regression network architecture directly predicts the 6 parameters used to describe an ellipse for tracking.

We also carried out video analysis of behavior in challenging environments including both 24-hour experimental videos that added bedding and a food cup to our open field arena, and videos from the open field experiment carried out as part of The Jackson Laboratory KOMP2 (Knockout Mouse Phenotyping Project)^8^ Phenotyping Center (Fig. 1a, column 5, 6, respectively). In the 24-hour experiment, we collected data over multiple days in which mice were housed in the open field with white paper bedding and food cup. The mice were kept in the open field in this multiday data collection paradigm, and continuous recording was carried out in light and dark conditions using an infrared light source. The bedding and food cups were moved by the mouse and the imaging light source alternated between infrared and visible light over the course of each day. The KOMP2 experiment uses a beam-break system in which mice are placed in a clear acrylic arena with infrared beams on all sides. Since the floor of the arena is clear acrylic, the surface of the table on which the arenas were placed shows through as dark grey. In addition, one arena was placed on the junction between two tables, leaving the joint visible. Further, the LED lights overhead caused a very high glare unique to each arena (Supplementary Video 2). This KOMP2 program has collected over five years of data using this system, and we wanted to carry out video-based recording as an added analysis modality to detect gait affects that cannot be identified by beam-break systems. Since environmental alterations could affect the behavioral output and legacy data interpretation, we could not optimize or otherwise alter the environment for video data collection. Instead, we simply added a camera on top of each arena. Traditionally, contrast and reflection hurdles could be overcome by changing the environment such that video data collection is optimized for analysis. For instance, to track albino mice, one can increase contrast by changing the background color of the open field to black. However, the color of the environment can effect the behavior of both mice and humans, and such manipulations can potentially confound the experimental results^3,4^. Regardless, such solutions will not work for piebald mice in a standard open field, or any mice in either the 24-hour data collection experiment or the KOMP2 arena.

We found that the combination of mouse coat colors and environments were difficult to handle with Ctrax (Supplementary Video 1) and LimeLight (Supplementary Video 3). We optimized and fine-tuned Ctrax for each video (Supplementary Methods) in each of the three experiments and still found a significant number frames with poor tracking performance (Fig. 1a, row 4). Such optimization or tuning of background model was not feasible with LimeLight. The frequency of poor tracking instances in an individual video increased as the environment became less ideal for tracking. Furthermore, the distribution of the errors was not random; for example, tracking was highly inaccurate when mice were in the corners, near walls, or on food cups (Fig. 1a, row 4), and less inaccurate when animals were in the center (Supplementary Video 1). While it is feasible to discard poorly tracked frames, this can lead to biased sampling and skewed biological interpretation.

We explored the cause of bad tracking across our three experiments and discovered that, in most cases, improper tracking was due to poor segmentation of the mouse from the background. This included both types of errors: Type I, instances when portions of the background are included as the foreground (e.g. shadows), and Type II, instances when portions of the mouse are removed from the foreground (e.g. albino mouse matching the background color). Since Ctrax uses a single background model algorithm, we tested whether other background model algorithms could improve tracking results. We tested 26 different segmentation algorithms^9^ and discovered that each of these traditional algorithms performs well under certain circumstances and fail in others (Fig. 1b). Other available tracking software packages including CADABRA^10^, EthoVision^11^, idTracker^12^, MiceProfiler^13^, MOTR^14^, Cleversys TopScan (http://cleversysinc.com/CleverSysInc/), Autotyping^15^, and Automated Rodent Tracker^16^, all of which rely on background subtraction approaches for tracking. Since all 26 background subtraction methods failed in some circumstances, we postulate that our results for Ctrax and LimeLight will hold true for these other technologies. In sum, although many video tracking solutions exist, none address the fundamental problem of mouse segmentation appropriately and generally rely on environmental optimization to achieve proper segmentation, therefore creating potential confounds with respect to robust data sampling and analysis. Thus, we could not overcome the fundamental issue of proper mouse segmentation in order to achieve high-fidelity mouse tracking with existing solutions.

A drawback in addition to the problem of inadequate mouse segmentation was the time cost for fine-tuning Ctrax’s settings or another background subtraction algorithm’s parameters. Fine-tuning the tracking settings for each video added significant time to our workflow when analyzing thousands of videos. For example, in tracking data from the 24-hour experiment, when mice were sleeping in one posture for an extended period of time, the mouse became part of the background model and could not be tracked. Typical supervision, such as using the Ctrax settings supervision protocol we outline in our methods, would take an experienced user 5 minutes of interaction for each hour of video to ensure high-quality tracking results. While this level of user interaction is tractable for smaller and more restricted experiments, large-scale and long-term experiments require a large time commitment to supervise the tracking performance.

We sought to overcome these difficulties by building a robust-next generation mouse tracker that uses neural networks and achieves high performance under complex and dynamic environmental conditions, is indifferent to coat color, and does not require persistent fine tuning by the user. Convolutional neural networks are computational models that are composed of multiple spatial processing layers that learn representations of data with multiple levels of abstraction. These methods have dramatically improved the state-of-the-art in speech recognition, visual object recognition, object detection, and many other domains such as drug discovery and genomics^17^. One of the key advantages of neural networks is that once an efficient network with suitable hyperparameters has been developed, it can easily be extended to other tasks by simply adding appropriate training data^18^. Thus, we sought to build a highly generalizable solution for mouse tracking.

We tested three primary neural network architectures for solving this visual tracking problem (Fig. 1d-e). Each approach attempted to describe the location of the animal through six variables: x and y location of the mouse in the matrix, major and minor axes of the mouse, and the angle the head is facing (Fig. 1c). To avoid the discontinuity of equivalent repeating angles, the networks predict the sine and cosine of the angle.

The first architecture is an encoder-decoder segmentation network that predicts a foreground-background segmented image from a given input frame (Fig. 1d). This network predicts on a pixel-wise basis whether there is a mouse or no mouse, with the output being a segmentation mask. The segmentation mask identifies all the pixels in the image that belong to the mouse. The primary structure of this architecture starts with a feature encoder, which abstracts the input image down into a small-spatial-resolution set of features. The encdoded features are then passed to a feature decoder that converts this set of features back into the same shape as the original input image. Additionally, the encoded features are also passed to three fully connected layers to predict which cardinal direction the ellipse is facing. We trained this feature decoder to produce a foreground-background segmented image. After the network produces this segmented image, we applied an ellipse-fitting algorithm for tracking (Supplementary Note 1).

The second network architecture is a binned classification network, whereby a probability distribution across a pre-defined range of possible values is predicted for each of the 6 ellipse-fit parameters (Fig. 1e). This network architecture begins with a feature encoder that abstracts the input image down into a small-spatial-resolution set of features. The encoded features are flattened and connected to additional fully connected layers whose output shape is determined by the desired resolution of the output. For instance, at a desired resolution of 1 pixel for the x-coordinate location of the mouse, there are 480 possible x-values to select from for a 480 x 480 px image. As such, the network contains 480 values (bins) to select from, one bin for each x-column in the 480 x 480 px image. When the network is run, the largest value in each heatmap is selected as the most probable value of the corresponding parameter. Each desired output parameter is realized as an independent set of trainable fully connected layers connected to the encoded features.

The third architecture is a regression network that predicts the numerical ellipse values directly from the input image (Fig. 1f). The network architecture begins with a feature encoder that abstracts the input down into a small spatial resolution. These encoded features are then flattened and connected to fully connected layers to produce an output shape of 6, the number of values that we ask the network to predict to fit an ellipse. We tested a variety of currently available general purpose feature encoders, and present data from the feature encoder Resnet V2 with 200 convolutional layers, which achieved the best performing results for this architecture^19^.

To test the neural network architectures, we built a training dataset of 16,234 training images and 568 separate validation images across multiple mouse strains and experimental setups (Supplementary Note 2). Annotated training images were augmented eightfold during training by applying reflections. Additionally, training images were further augmented by adding small random changes in contrast, brightness, and rotations to make the network robust to minor fluctuations in input data. We created an OpenCV-based labeling interface for creating our training data (Supplementary Methods) that allows us to quickly label foreground and background, and fit an ellipse (Supplementary Fig. S1). This labeling interface can be used to quickly generate annotated training data in order to adapt any network to new experimental conditions through transfer learning.

Our network architectures were built, trained, and tested in Tensorflow v1.0, an open-source software library for designing applications that use neural networks^20^. Training benchmarks presented were conducted on the Nvidia P100 GPU architecture. We tuned the hyperparameters through several training iterations. After the first training of networks, it was observed that the networks performed poorly under particular circumstances that had not been included in the annotated data, including mid-jump, odd postures, and urination in the arena. We identified and incorporated these difficult frames into our training dataset to further improve performance. A full description of the network architecture definitions and training hyperparameters are available (Supplementary Methods, Supplementary Table I). Overall, training and validation loss curves indicated that each of the three network architectures trains to a performance with an average error between 1 and 2 pixels (Fig. 2a). The encoder-decoder segmentation architecture converged to a validation error of 0.9px (Fig. 2 a, b, c). Surprisingly, upon inspection of the validation curve for the binned classification network we found that it displayed unstable loss curves, indicating overfitting and poor generalization (Fig. 2b, e). The regression architecture converged to a validation error of 1.2 px, showing a better training than validation performance (Fig. 2a, b, d).

**Figure 2:**
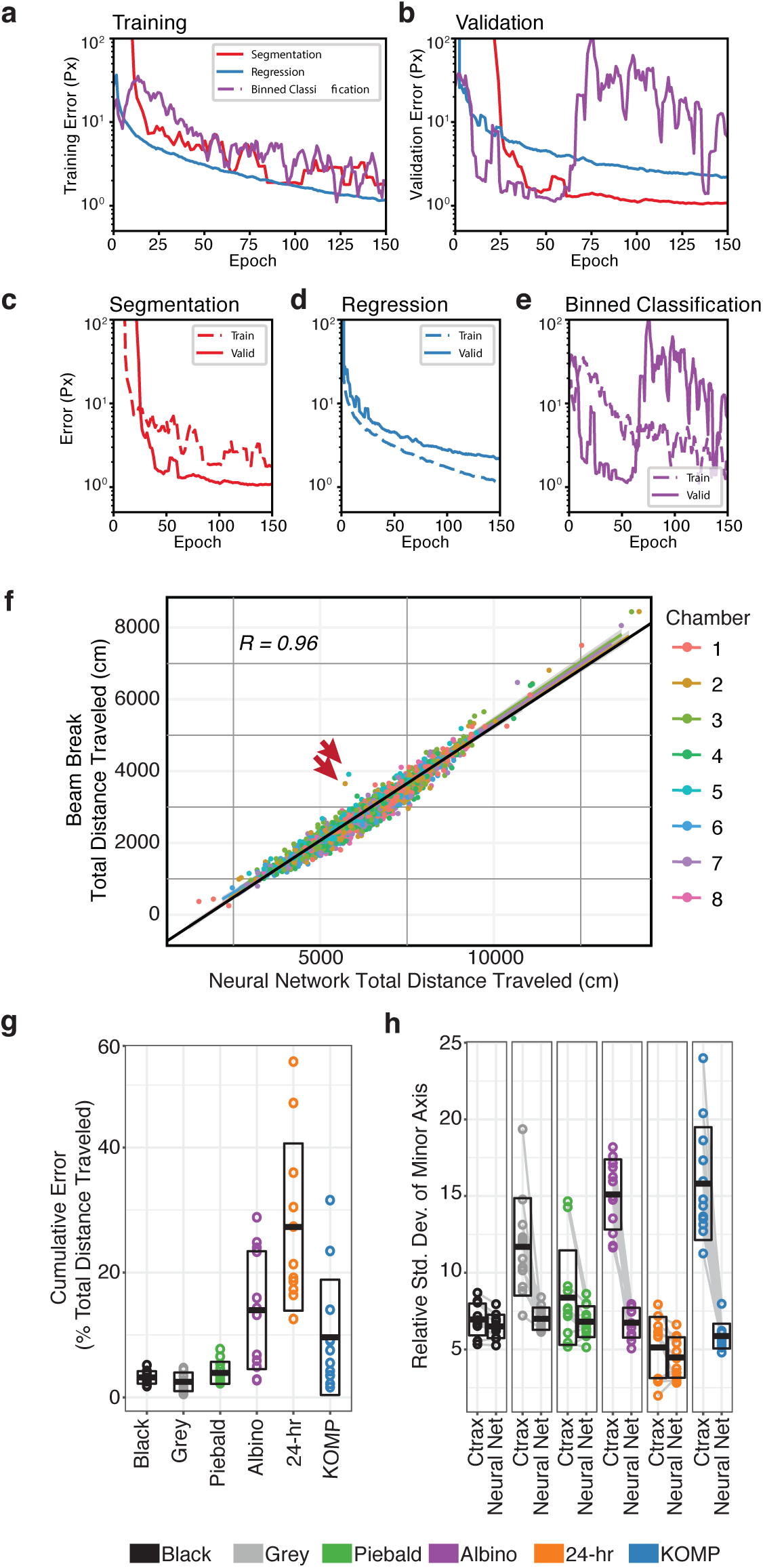
(**a-e**) Performance of our tested network architectures during trainings. (**a**) Training curves show comparable performances of the three architectures during training, independent of the network architecture. (**b**) Validation curves show different performances across the three network architectures. The encoder-decoder segmentation network performs the best. (**c, d, e**) Comparison of training and validation performance curves, by network architecture type. (**c**) Performance increases for validation in our encoder-decoder segmentation network architecture. (**d**) Performance decreases for validation in our regression network architecture, but a good generalization performance is maintained by asymptotically converging to a value. (**e**) The binned classification network architecture becomes unstable at 55 epochs of training, even though the training curve shows continued improved performance at this timepoint. (**f**) Comparing our encoder-decoder segmentation network architecture with a beam break system, we observe a high correlation. Each point represents an individual video tracked using both our neural network and a beam break system. Our network performs consistently, even though the arenas are visually different from one another. We identify two videos of individual mice that deviate the from this trend (red arrows). (**g**) Predictions from two approaches yield high agreement on environments with high contrast between the mouse and background (black, grey, and piebald mice in the white background open-field assay). As the segmentation problem becomes more computationally difficult, the relative error increases (albino mice in the white background open-field assay black mice in the 24-hr assay, KOMP2 experiment). Due to low activity in the 24-hr setup, minor errors in tracking have a large influence on measurements of thetotal distance traveled. Points indicate individuals in a group, bars indicate mean +/- standard deviation. (**h**) Relative standard deviation of the minor axis maintains a high correlation when the mouse and environment have a high contrast (black mice in the white background open-field assay). When segmentation includes shadows, includes reflections, or removes portions of the mouse, the minor axis length is not properly predicted and increases the relative standard deviation (grey, piebald, and albino mice in the white background open-field assay, black mice in the 24-hour assay, KOMP2 experiment). Points indicate individuals in a group, bars indicate mean +/- standard deviation.

Not only does the encoder-decoder segmentation architecture perform well, but it also is computationally efficient for GPU compute, requiring an average processing time of 5-6ms per frame. With the encoder-decoder segmentation architecture, our video data could be processed at a rate of up to 200 frames per second (fps) (6.7X realtime) on a Nvidia P100, which is a server-grade GPU.; and a rate of up to 125 fps (4.2X realtime) on a Nvidia TitanXP, a consumer-grade GPU. This high processing speed is likely due to the structure of the encoder-decoder segmentation architecture, as it is only 18 layers deep and contains only 10.6 million trainable parameters. In comparison, Resnet V2 200, the feature extractor that gave the best results for the regression architecture, is a large and deep network with over 200 layers and 62.7 million trainable parameters and leads to a substantially longer processing time per frame (33.6ms on a Nvidia P100). Other pre-built general-purpose networks^21^ achieve similar or worse performances at a tradeoff of faster compute time (data not shown). Thus, regression networks are an accurate but computationally expensive solution.

We also tested the minimum training dataset size required to train the encoder-decoder segmentation network, by randomly subsetting our training dataset to smaller numbers of annotated images (10,000 to 500) and training the network from the beginning. Surprisingly, we obtained good results from a network trained with only 2,500 annotated images, a task that takes approximately three hours to generate with our labeling interface (Supplementary Fig. S2). Given the computational efficiency, accuracy, and training stability of the encoder-decoder segmentation architecture, and the small training dataset size that it requires, we concluded that this architecture is optimal for our needs. We used this trained neural network to predict the location of mice for entire videos and compare tracking performance with other non-neural network approaches including a beam-break system (KOMP2) and a video tracking system (Ctrax).

We evaluated the quality of the encoder-decoder segmentation neural network tracking architecture by inferring entire videos from mice with disparate coat colors and data collection environments (Fig. 1a) and visually evaluating the quality of the tracking. We also compared this neural network-based tracking architecture with an independent modality of tracking, the KOMP2 beam-break system (Fig. 1a, column 6). We tracked 2,002 videos of individual mice comprising 700 hours of video from the KOMP2 experiment using the encoder-decoder segmentation neural network architecture and compared the results with the tracking data obtained using the KOMP2 beam-break system (Fig. 2f). These data comprised mice of 232 knockout lines on the C57BL/6NJ background that were tested in 20-minute open field assay in 2016 and 2017. Since each KOMP2 arena has slightly different background due to the transparent and reflective walls, we compared tracking performances of the two approaches for each of the eight testing arenas used in the 2016 and 2017 KOMP2 open-field assays (Fig. 2f, colors shows arena), and compared tracking performances for all the arenas combined (Fig. 2f, black line). We observed a very high correlation between the total distance traveled in the open field as measured by the two approaches across all eight KOMP2 testing arenas (R = 96.9%, Fig. 2f). We observed two animals with high discordance from this trend (Fig. 2f, red arrows). Observation of the video showed odd behaviors for both animals, with a waddle gait in one and a hunched posture in the other (Supplementary Video 2). We postulate that these behaviors led to abnormal beams breaks causing erroneously high total distances traveled measured via the beam break system. This example highlights an important advantage of the neural network, as it is unaffected by the behavior of the animal.

We then compared the performance of our trained segmentation neural network with the performance of Ctrax across a broad selection of videos from the various testing environments and coat colors previously tracked using Ctrax and LimeLight (Fig. 1a). We wish to emphasize that we compared the performance of our network with that of Ctrax because Ctrax is one of the best conventional tracking software packages that allows fine tuning of the many tracking settings, is open source, and provides user support. Given the results with the 26 background subtraction approaches (Fig. 1b), we expected similar or worse performances from other tracking systems. We tracked 72 videos, broken into 6 groups (Fig. 1a) with 12 animals per group, with both our trained encoder-decoder segmentation neural network and Ctrax. The settings for Ctrax were fine-tuned for each of the 72 videos, as described in ‘Ctrax Settings Supervision Protocol’ in Supplementary Methods. Videos from the 24-hr experiment showing that animals that were sleeping continually for the full video duration (one hour) were manually omitted from comparison, as Ctrax will incorporate the mouse as part of the background model. We calculated a cumulative relative error of total distance traveled between Ctrax and our neural network (Fig. 2g). Specifically, for every minute in the video, we compared the distance-traveled prediction of the neural network with that of Ctrax. This metric measures the accuracy of center of mass tracking of each mouse. Tracking for black, gray, and piebald mice in the white-background open-field apparatus showed errors less than 4%; however, significantly higher levels of error were seen in albino mice in the open-field arena with a white floor (14%), black mice in the 24-hour arena (27%), and black mice in the KOMP2 testing arena (10%) (Fig. 2g and Supplementary Video 1). Thus, we could not adequately track albino mice in the open-field arena with a white floor, black mice in the 24-hour arena, or black mice in the KOMP2 testing arena without the neural network tracker.

We also observed, using Ctrax, that when foreground segmentation prediction is incorrect, such as when shadows are included in the prediction, the ellipse fit does not correctly represent the posture of the mouse (Supplementary Video 1). In these cases, even though the center of mass tracking was acceptable, the ellipse fit itself was highly variable. Modern machine learning software for behavior recognition, such as the Janelia Automatic Animal Behavior Annotator (JAABA)^22^, utilize the time series of ellipse fit tracking for classification of behaviors. We quantitated the stability of ellipse tracking through measuring the relative standard deviation of the minor axis and comparing approaches. This metric shows the least variance across all sizes of laboratory mice, as the width of an individual mouse remains similar through a wide range of postures expressed in behavioral assays when tracking is accurate. We observed a high level of tracking variation with grey and piebald mice in the white open field arena (Fig. 2h) even though there is low cumulative relative error of total distance traveled (Fig. 2g). As expected, we observed a high relative standard deviation of the minor axis for albino mice (white open field arena) and KOMP2 tracking. Thus, for both center of mass tracking and variance of ellipse fit we find that the neural network tracker outperforms traditional background subtraction-based trackers.

Having established the encoder-decoder segmentation neural network as a highly accurate tracker, we tested its performance using two large behavioral experiments. For the first experiment, we generated white-surfaced open-field video data with 1,845 mice, including 58 strains of mice including mice with diverse coat colors, piebald mice, nude mice, and obese mice; and covering a total of 1,691 hours (Fig. 3a). This dataset consists of 47 inbred strains and 11 isogenic F1 strains and is the largest open-field dataset generated, based on the data in the Mouse Phenome Database^23^. Using a single trained network without any user tuning, we were able to track all mice with high accuracy. We visually checked mice from a majority of the strains for fidelity of tracking and observed excellent performance. The activity phenotypes that we observed agree with previously published datasets of mouse open-field behavior^23^. For the second dataset, we tracked 24-hour video data collected for four C57BL/6J and two BTBR T^+^ ltpr3^tf^/J mice (Fig. 1a, column 5). These mice were housed with beddingand a food cup over multiple days during which the food changed location and under 12:12 light-dark conditions. Video data were recoded using visible and infrared light sources. We tracked activity across all animals under these conditions using the same encoder-decoder segmentation neural network architecture used for the first experiment, and observed very good performance under light and dark conditions (Fig. 3b, light and dark blue points, respectively). As expected, we observed daily activity rhythm with high levels of locomotor activity during the dark phase (Fig. 3b, red curve).

**Figure 3:**
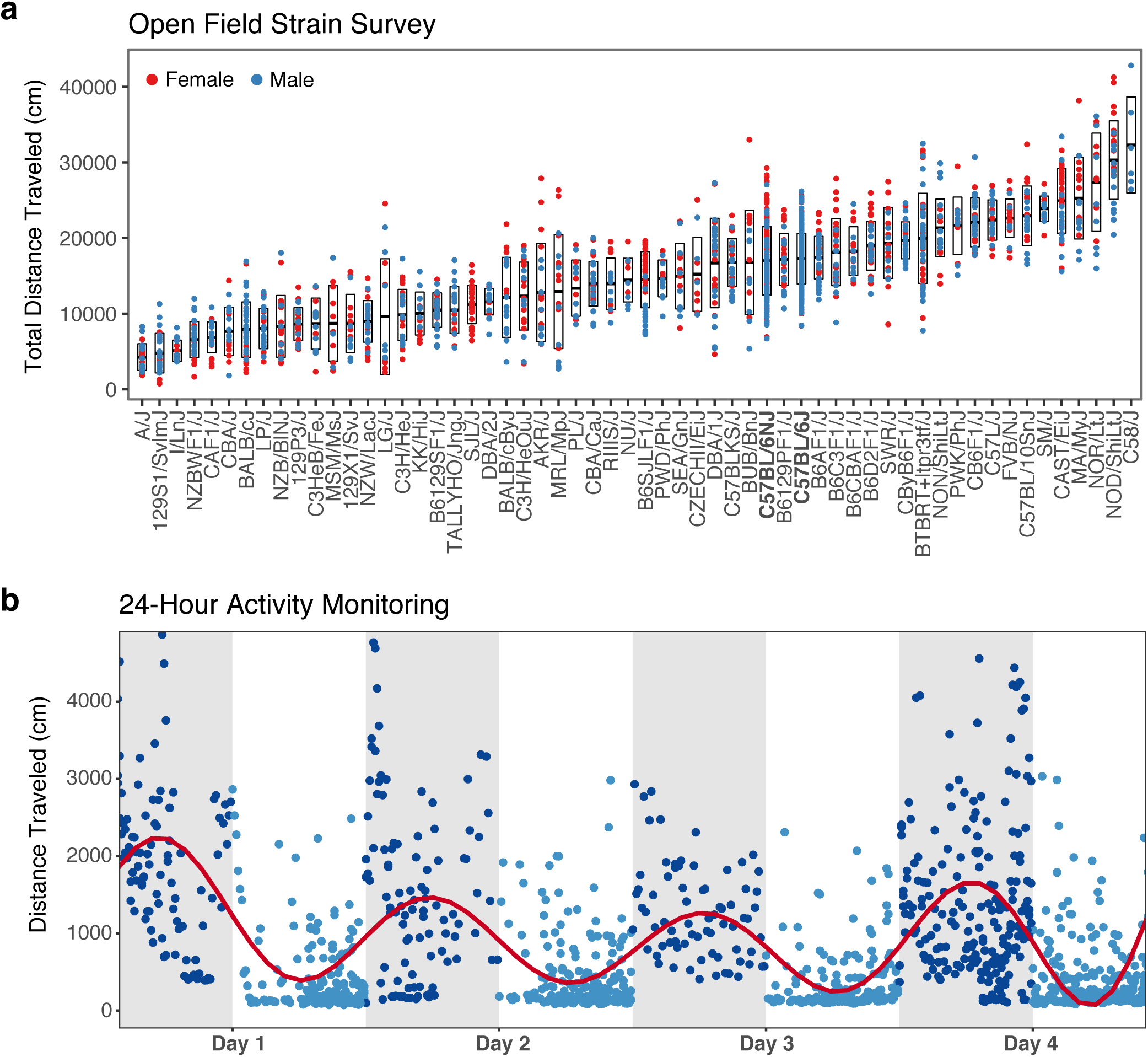
Highly scalable tracking with a single neural network. (**a**) A large strain survey showing genetically diverse animals traced with our encoder-decoder segmentation network. 1,845 animals including 58 inbred and F1 isogenic strains, totaling 1,691 hours of video, were processed by a single trained neural network without any user-involved fine-tuning. Total distance traveled in a 55-minute open field assay is shown. Points indicate individuals in a strain, bars indicates mean +/- standard deviation. Two reference mouse strains are shown in bold, C57BL/6J and C57BL/6NJ (**b**) Daily activity rhythms were observed in six animals continuously tracked over 4 days in a dynamic environment with our encoder-decoder segmentation neural network. Points indicate distance traveled in an epoch. Red line indicates polynomial fit showing

Video-based tracking of animals in complex environments has been a long-standing challenge in the field of animal behavior^24^. Current state-of-the-art animal-tracking systems do not address the fundamental issue of animal segmentation and rely heavily on visual contrast between the foreground and background for accurate tracking. As a result, the user must restrict the environment to achieve optimal results. Here we describe a modern neural network-based tracker that is able to function in complex and dynamic environments. Our network addresses a fundamental issue in tracking—foreground and background segmentation—by using a trainable neural network. We test three different architectures and find that an encoder-decoder segmentation network architecture achieves the highest level of accuracy and functions at a high speed (over 6X real time). Furthermore, we provide a labeling interface that allows the user to train a new network for their specific environment by labeling as few as 2,500 images, which takes approximately 3 hours. We compare our network to two existing solutions and find that it vastly outperforms them in complex environments. We expect similar results with any off-the-shelf system that utilizes traditional background subtraction approaches. In fact, when we tested 26 different background subtraction methods we discovered that each failed under certain circumstances. However, a single neural network architecture functions for all coat colors of mice under multiple environments without the need for fine tuning or user input. Our machine learning approach enables long-term tracking under dynamic environmental conditions with minimal user input, thus establishing the basis of the next generation of tracking architecture for behavioral research.

## Methods

Methods, including statements of data availability, associated code, and details of experimental setups, are available in the Supplementary Methods section.

## Acknowledgments

We thank the members of the Kumar laboratory for suggestions and editing of the manuscript. We thank JAX Information Technology team members Edwardo Zaborowski, Shane Sanders, Rich Brey, David McKenzie, and Jason Macklin for infrastructure support; and we thank KOMP2 behavioral testers James Clark, Pamelia Fraungruber, Rose Presby, Zachery Seavey, and Catherine Witmeyer. This work used the National Science Foundation (NSF) Extreme Science and Engineering Discovery Environment (XSEDE) XStream service at Stanford University through allocation TG-DBS170004. Funding also included the NIH grant DA041668 from NIDA, a Brain and Behavioral Foundation Young Investigator Award, and a JAX Director’s Innovation Fund to V. K. We thank Nvidia GPU grant program for providing a TitanXP for our work.

## Author Contributions

V. K. and B. Q. G conceived the project and analyzed the data. B. Q. G and K. J. F. performed the computational work, S. P. D. and B. Q. G. carried out behavior tests, and S. A. M., R. E. B., J. K. W., E. J. C., and C. M. L. planned and analyzed strain survey and KOMP2 data. B. Q. G and V. K. wrote the manuscript and designed the figures.

## Online Methods

### Experimental Arenas

#### Open Field Arena

Our open field arena measures 52cm by 52cm by 23cm. The floor is white PVC plastic and the walls are grey PVC plastic. To aid in cleaning maintenance, a white 2.54cm chamfer was added to all the inner edges. Illumination is provided by an LED ring light (Model: F&V R300). The ring light was calibrated to produce 600 lux of light in each of our 24 arenas.

#### 24-Hour Monitoring Open Field Arena

We augmented 6 of our open field arenas for multiple day testing. We set our overhead LED lighting to a standard 12:12 light-dark cycle. ALPHA-dri was placed into the arena for bedding. To provide food and water, a single Diet Gel 76A food cup was placed in the arena. This nutritional source was monitored and replaced when depleted. Each arena was illuminated at 250 lux during the day and <5 lux during the night. For recording videos during the night, additional IR LED (940nm) lighting was added.

#### KOMP2 Open Field Arena

In addition to our custom arenas, we also benchmarked our approach on a commercially available system. The Accuscan Versamax Activity Monitoring Cages is constructed using clear plastic walls. As such, visual tracking becomes very difficult due to the consequent reflections. The cage measures 42cm by 42cm by 31cm. Lighting for this arena was via LED illumination at 100-200 lux.

### Video Acquisition

#### Imaging Hardware

All data was acquired using the same imaging equipment. Data was acquired at 640×480px resolution, 8-bit monochrome depth, and 30fps using Sentech cameras (Model: STC-MB33USB) and Computar lenses (Model: T3Z2910CS-IR). Exposure time and gain were controlled digitally using a target brightness of 190/255. Aperture was adjusted to its widest so that lower analog gains were used to achieve the target brightness. This in turn reduced amplification of baseline noise. Files were saved temporarily on a local hard drive using the “raw video” codec and “pal8” pixel format. Our typical assays run for two hours, yielding a raw video file of approximately 50GB. Overnight, we use FFmpeg software (https://www.ffmpeg.org/) to apply a 480×480px crop, de-noise filter, and compress using the mpeg4 codec (quality set to max), which yields a compressed video size of approximately 600MB.

One camera and lens was mounted approximately 100cm above each arena to alleviate perspective distortion. Zoom and focus were set manually to achieve a zoom of 8px/cm. This resolution both minimizes the unused pixels on our arena border and yields approximately 800 pixels area per mouse. Although the KOMP2 arena is slightly smaller, the same zoom of 8px/cm target was utilized.

### Ctrax Settings Supervision Protocol

Ctrax contains a variety of settings to enable optimization of tracking^1^. The authors of this software strongly recommend, first and formost, ensuring that he arena is set up under specific criteria to ensure good tracking. In most of our tests, we intentionally use an environment in which Ctrax is not designed to perform well (e.g., albino mice on a white background). That being said, with well-tuned parameters, a good performance is still achievable. However, with a large number of settings to manipulate, Ctrax can easily require substantial time to achieve a good tracking performance. Here, we describe our protocol for setting up Ctrax for tracking mice in our environments.

First, we create a background model. The core of Ctrax is based on background subtraction, and thus a robust background model is essential for functionality. Models function optimally when the mouse is moving. To create the background model, we seek to a segment of the video in which the mouse is clearly moving, and we sample frames from that section. This ensures that the mouse is not included in the background model. This approach significantly improves Ctrax’s tracking performance on our 24-hour data, as the mouse moves infrequently due to sleeping and would typically be incorporated into the background model.

The second step is to set the settings for background subtraction. Here, we use the Background Brightness normalization method with a Std Range of 254.9 to 255.0. The thresholds applied to segment out the mouse are tuned on a per-video basis, as slight changes in exposure and coat color will influence the performance. To fine-tune these thresholds, we apply starting values based on previous videos analyzed and adjust values by checking multiple portions of the video. Every video is inspected for proper segmentation on difficult frames, such as the mouse rearing on the wall. Additionally, we apply morphological filtering to both remove minor noise in the environment as well as remove the tails of mice for fitting an ellipse. We use an opening radius of 4 and a closing radius of 5.

Lastly, we manually set a variety of tracking parameters that Ctrax enables to ensure that the observations are in fact mice. For optimal time efficiency, these parameters were tuned well once and then used for all other mice tracked. If a video was performing noticeably poorly, the general settings were tweaked to improve performance. For the shape parameters, we computed bounds based on two standard deviations from an individual black mouse video. We lowered the minimum values further because we expected that certain mice would perform poorly on the segmentation step. This allows Ctrax to still find a good location of the mouse despite not being able to segment the entire mouse. This approach functions well, as all of our setups have the same zoom of 8, and the mice tested are generally the same shape. Motion settings are very lenient, because our experimental setup tracks only one mouse in the arena at a time. Under the observation parameters, we primarily utilize the “Min Area Ignore” setting to filter out detections larger than 2,500 pixels. Under the hindsight tab, we use the “Fix Spurious Detections” setting to remove detections with a length shorter than 500 frames.

### Training Sets

#### Labeling Software

We annotated our own training data using custom software that was written to accommodate obtaining the necessary labels. We used the OpenCV library (https://opencv.org/) to create an interactive watershed-based segmentation and contour-based ellipse-fit. Using the software GUI we developed, the user left-clicks to mark points as the foreground (a mouse) and right-clicks to label other points as the background (Supplementary Fig. 1). Upon a keystroke, the watershed algorithm is executed to predict a segmentation and ellipse. If users need to make edits to the predicted segmentation and ellipse, they can simply mark additional areas and run the watershed again. When the predictions are of sufficiently high quality, users then select the direction of the ellipse. They do this by selecting one of four cardinal directions: up, down, left, right. Since the exact angle is selected by the ellipse-fitting algorithm, users need only to identify the direction ±90 degrees. Once a direction is selected, all the relevant data is saved to disk and users are presented with a new frame to label. Full details on the software controls can be found in the software documentation.

The objective of our annotated dataset is to identify good ellipse-fit tracking data for mice. While labeling data, we optimized the ellipse-fit such that the ellipse was centered on the mouse’s torso with the major axis edge approximately touching the nose of the mouse. Frequently, the tail was removed from the segmentation mask to provide a better ellipse-fit. For training networks for inference, we created three annotated training sets. Each training dataset includes a reference frame (input), segmentation mask, and ellipse-fit. Each training set was generated to track mice in a different environmental setup.

### Neural Network Models

The neural networks we trained fall into three categories: segmentation, regression, and binning. Our tested models can be viewed visually in Fig. 1d-f.

The first network architecture is modeled after a typical encoder-decoder structure for segmentation (Fig. 1d). The first half of the network (encoder) utilizes 2D convolutional layers followed by batch normalization, a ReLu activation, and 2D max pooling layers. We use a starting filter size of 8 that doubles after every pooling layer. The kernels used are of shape 5×5 for 2D convolution layers and 2×2 for max pooling layers. Our input is of shape 480×480×1 and after six of these repeated layers, the resulting shape is 15×15×128. We apply another 2D convolutional layer (kernel 5×5, 2x filters) followed by a 2D max pool with a different kernel of 3×3 and stride of 3. One final 2D convolutional layer is applied to yield our feature bottleneck with a shape of 5×5×512. This feature bottleneck is then passed to both the segmentation decoder and angle predictor. The segmentation decoder reverses the encoder using strided transpose 2D convolutional layers and carries over pre-downsampled activations through summation junctions. It should be noted that this decoder does not utilize ReLu activations. After the layers return to the 480×480×8 shape, we apply one additional convolution, with a kernel size of 1×1, to merge the depth into two images: background prediction and foreground prediction. We apply a softmax function across this depth. From the feature bottleneck, we also create a prediction for angle prediction. We achieve this by applying two 2D convolution layers with batch normalization and ReLu activations (kernel size 5×5, feature depths 128 and 64). From here, we flatten and use one fully connected layer to yield a shape of four neurons, which function to predict the quadrant in which the mouse’s head is facing. Since the angle is predicted by the mask, we need only to select the correct direction (± 180 deg). The four possible directions that the network can select are 45-135, 135-225, 225-315 and 315-45 degrees on a polar coordinate grid. These boundaries were selected to avoid discontinuities in angle prediction.

The second network architecture is a binned regression approach (Fig. 1e). Instead of predicting the parameters directly, the network instead selects the most probable value from a selection of binned possible values. The major difference between this structure and a regression structure is that the binned regression network training relies on a cross entropy loss function whereas a regression network relies on a mean squared error loss function. Due to memory limitations, we tested only custom VGG-like networks with reduced feature dimensions. Our best-performing network is structured with two 2D convolutional layers followed by a 2D max pooling layer. The kernels used are of shape 3×3 for 2D convolutional layers and 2×2 for 2D max pooling layers. We start with a filter depth of 16 and double after every 2D max pool layer. This two convolutional plus max pool sequence is repeated five times to yield a shape of 15×15×256. This layer is flattened and connected to a fully connected layer for each output ellipse-fit parameter. The shape of each output is dictated by the desired resolution and range of the prediction. For testing purposes, we observed only the center location and trained with a range of the entire image (0-480). Additional outputs, such as angle prediction, could simply be added as additional output vectors.

The third network architecture is modeled after a typical regression predictor structure (Fig. 1f). While the majority of regression predictors realize the solution through a bounding box, an ellipse simply adds one additional parameter: the angle of the mouse’s head direction. Since the angle is a repeating series with equivalence at 360deg and 0deg, we transform the angle parameter into its sine and cosine components. This yields a total of six parameters regressed from the network. The first half of this network encodes a set of features relevant to correctly predicting the six parameters. From the encoded feature set, we flatten the network and applied a fully convolutional layer to regress the parameters for the ellipse-fit. We tested a wide variety of pre-built feature detectors including Resnet V2 50, Resnet V2 101, Resnet V2 200, Inception V3, Inception V4, VGG, and Alexnet. In addition to these pre-built feature detectors, we also surveyed a wide array of custom networks. Of these general purpose feature encoders and custom networks, Resnet V2 200 performed the best.

### Neural Network Training

This section describes all of the procedures pertaining to training our neural network models. The three procedures described here are training set augmentation, training hyperparameters, and a benchmark for training set size.

Training set augmentation has been an important aspect of training neural networks since Alexnet^2^. We utilize a handful of training set augmentation approaches to achieve good regularization performance. Since our data is from a birds-eye view, it is straightforward to apply horizontal, vertical, and diagonal reflections for an immediate 8x increase in our equivalent training set size. Additionally, at runtime, we apply small rotations and translations for the entire frame. Rotation augmentation values are sampled from a uniform distribution. Finally, we apply noise, brightness, and contrast augmentations to the frame. The random values used for these augmentations are selected from a normal distribution.

Hyperparameters, such as training learn rate and batch size, were selected independently for each network architecture trained. While larger networks, such as Resnet V2 200, can run into memory limitations for batch sizes at an input size of 480×480, good learn rate and batch size were experimentally identified using a grid search approach^3^. Supplementary Table I summarizes all the hyperparameters selected for training these network architectures.

We also benchmarked the influence of training set size on network generalization in order to determine the approximate amount of annotated training data required for good network performance of the encoder-decoder segmentation network architecture (Supplementary Fig. 2). We tested this benchmark by shuffling and randomly sampling a subset of the training set. Each subsampled training set was trained and compared to an identical validation set. While the training curves appear indistinguishable, the validation curves tained with fewer than 2,500 training annotations diverge from the group. This suggests that the training set is no longer large enough to allow the network to generalize well. While the exact number of training samples will ultimately rely on the difficulty of the visual problem, a recommended starting point would be around 2,500 training annotations.

### Animals Used

All animals were obtained from The Jackson Laboratory production colonies. Adult mice aged 8 to 14 weeks were behaviorally tested in accordance with approved protocols from The Jackson Laboratory Institutional Animal Care and Use Committee guidelines. Open field behavioral assays were carried out as previously described^4^. Briefly, group-housed mice were weighed and allowed to acclimate in the testing room for 30-45 minutes before the start of video recording. Data from the first 55 minutes of activity are presented here. Where available, 8 males and 8 females were tested from each inbred strain and F1 isogenic strain.

### Code and Training Set Availability

Neural network training and inference code as well as annotated datasets will become available upon publication.

## Supplementary Note 1

### Fitting an Ellipse to a Mask

The same ellipse-fit algorithm was used as described in supplemental section 4.4.2 of the Ctrax paper^1^. While the paper uses a weighted sample mean and variance for these calculations, the segmentation neural network retains invariance to the situations in which they describe improvements. Additionally, we observe no significant difference between using weighted and unweighted sample means and variances.

Given a segmentation mask, the sample mean of pixel locations is calculated to represent the center position.

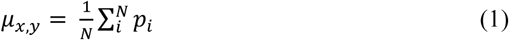

Similarly, the sample variance of pixel locations is calculated to represent the major axis length (*a*), minor axis length (*b*), and angle (θ).

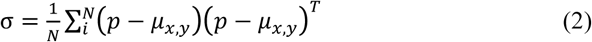

To obtain the axis lengths and angle, an eigenvalue decomposition equation must be solved.

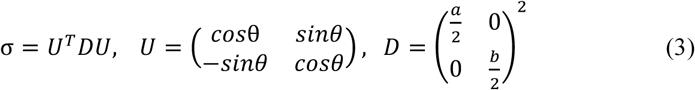

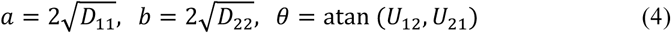

## Supplementary Note 2

### Annotated Datasets

We created 3 annotated datasets for training neural networks, each including a reference frame (input), segmentation mask, and ellipse-fit. Each dataset was generated to track mice in a different environment.

The first annotated dataset uses images sampled from our standard open field arena video experiment and contains 16,802 annotated frames. The first 16,000 frames were annotated from 65 separate videos acquired from one of 24 testing arenas. An additional 802 frames across 50 additional videos that perform poorly were identified, correctly annotated, and incorporated into the annotated dataset to further generalize and improve performance.

The second annotated dataset uses images sampled from our 24-hour experiment, which uses the standard open field arena with ALPHA-dri bedding and a food cup under two distinct lighting conditions (day visible illumination and night infrared illumination). For the dataset from this environment, we annotated a total of 2,192 frames across 6 videos of 4 day duration. Of the total number of annotated frames, 916 were taken from night illumination and 1,276 from the daylight illumination.

The third annotated dataset uses images sampled from the Accuscan Versamax Activity Monitoring Cages for the KOMP2 experiment. The dataset for this environment comprised 1,083 annotated frames. These annotations were all sampled across different videos (1 frame labeled per video) and 8 different arenas.

**Supplementary Table I.** Training Hyperparameters

Training Hyperparameters
| Model | Parameter | Value |
| --- | --- | --- |
| All three | Rotation | $\pm 2.5$ deg |
| | Translation | $\pm 5.0$ px |
| | Additive Noise | $\mu=0.0$, $\sigma=5.0$ |
| | Brightness | $\pm 5\%$ |
| | Contrast | $\pm 5\%$ |
| Encoder-Decoder Segmentation Network | Learning Rate | $10^{-5}$ |
|  | Batch Size | 50 |
| Regression Network | Learning Rate | $10^{-5}$ |
|  | Batch Size | 5 |
| Binned Classification Network | Learning Rate | $10^{-3}$ |
|  | Batch Size | 25 |

**Supplementary Figure S1:**
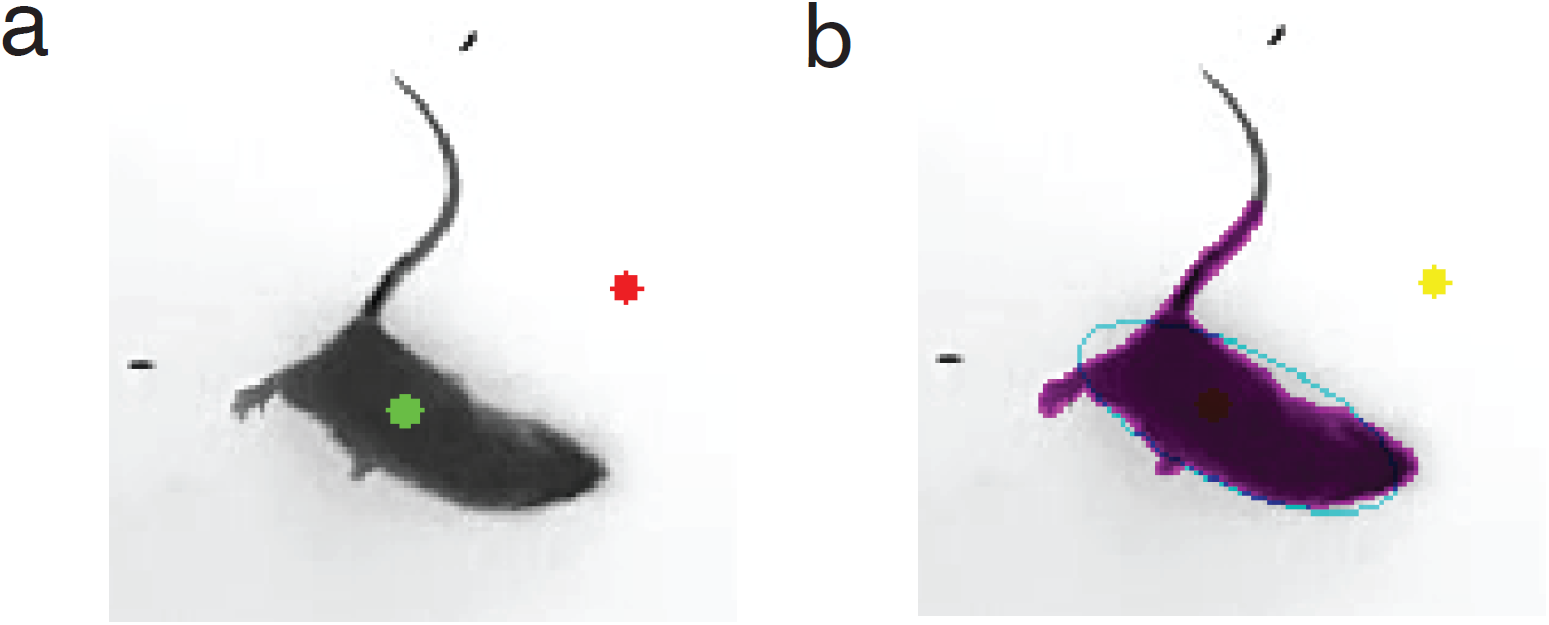
An example of our labeling GUI software. (**a**) shows that the user has zoomed into the mouse and placed two marks: one for foreground (green), and one for background (red). (**b**) shows the resulting segmentation (magenta), ellipse-fit (cyan), and old background (yellow) annotations.

**Supplementary Figure S2:**
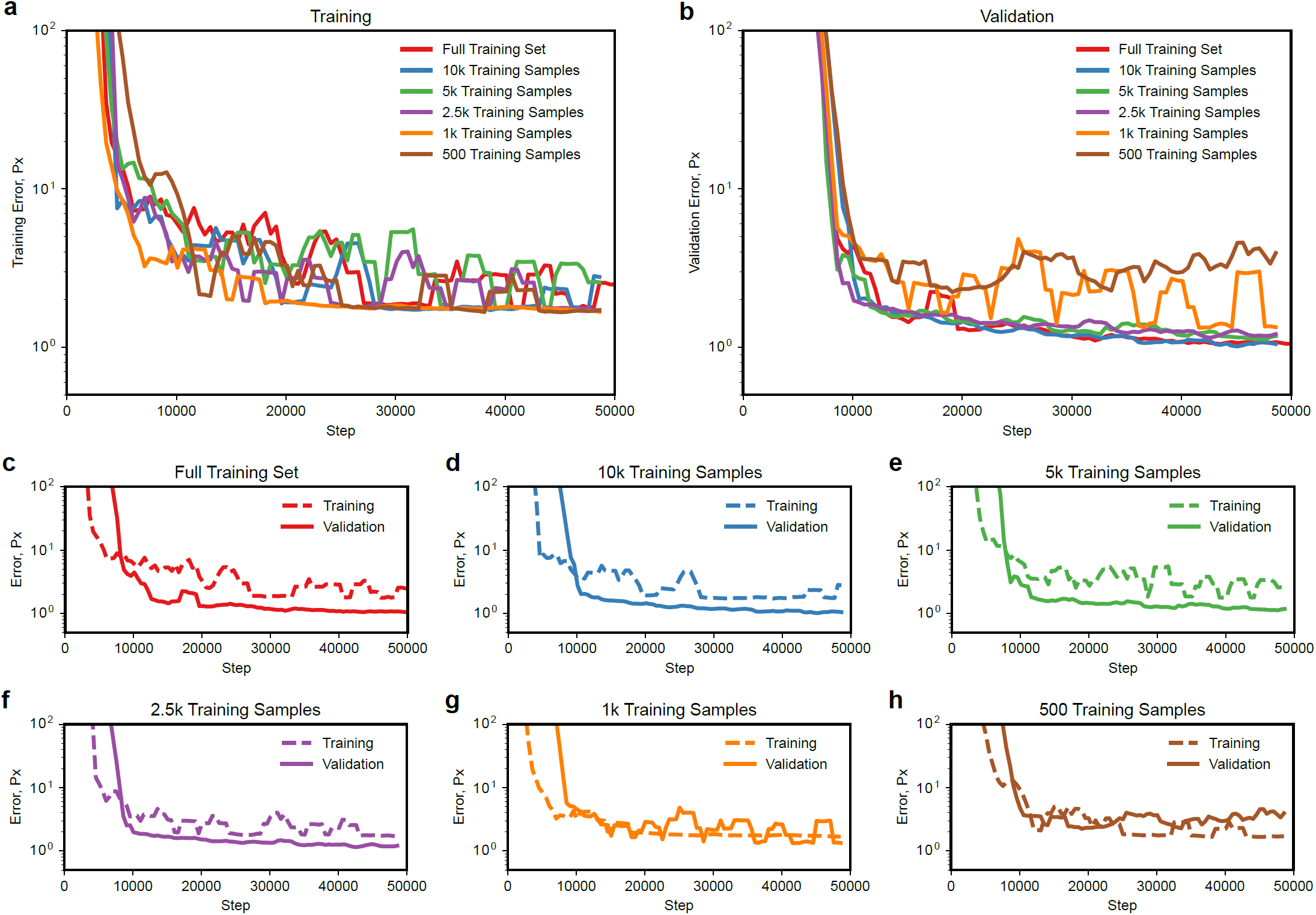
We benchmarked how the training-set size influences the performance of a trained encoder-decoder segmentation network. Full training set includes 16,234 annotated frames. (**a**) Training-set size does not impact training set error rate. (**b**) Validation performance converges to the same value above 2,500 training samples, but the error rate increases when 1,000 or fewer training samples are used. (**c-f**) Validation accuracy outperforms training accuracy when 2,500 or more training samples are used. (**g**) Validation accuracy begins to show signs of weak generalization by only matching, and not exceeding, training accuracy at 1,000 training samples. (**h**) A network trained using only 500 training samples is clearly overtraining, shown by the diverging and increasing validation error rate.

